# The Existence of At Least Three Genomic Signature Patterns and At Least Seven Subtypes of COVID-19 and the End of the Disease

**DOI:** 10.1101/2022.01.24.477579

**Authors:** Zhengjun Zhang

## Abstract

Hoping to find genomic clues linked to COVID-19 and end the pandemic has driven scientists’ tremendous efforts to try all kinds of research. Signs of progress have been achieved but are still limited. This paper intends to prove the existence of at least three genomic signature patterns and at least seven subtypes of COVID-19 driven by five critical genes (the smallest subset of genes). These signatures and subtypes provide crucial genomic information in COVID-19 diagnosis (including ICU patients), research focuses, and treatment methods. Unlike existing approaches focused on gene fold-changes and pathways, gene-gene nonlinear and competing interactions are the driving forces in finding the signature patterns and subtypes. Furthermore, the method leads to 100% accuracy, which shows biological and mathematical equivalences between COVID-19 status and the signature patterns and a methodological advantage over other existing methods that cannot lead to 100% accuracy. As a result, as new biomarkers, the new findings can be much more informative than other findings for interpreting biological mechanisms, developing the second (third) generation of vaccines, antiviral drugs, and treatment methods, and eventually bringing new hopes to an end of the pandemic.

## 1 Introduction

Since the virus SARS-COV-2 was first reported in January 2020, numerous research results related to the virus and COVID-19 disease have been published. Scientists have put tremendous effort into trying to find genomic clues linked to COVID-19. However, knowledge of COVID-19 is still limited, and many problems have remained unanswered^1;2;3;4;5;6;7;8;9;10;11;12^. Many published results cannot guarantee 100% accuracy. With an exception, a data science study^2^ reported five critical genes and their combined effects can 100% accurately classify COVID-19 patients and COVID-19 free patients into their respective groups and further classify COVID-19 patients into seven subtypes.

The 100% accuracy enables scientists to focus on and further explore the biological mechanisms among COVID-19 specific genes (note that many reported genes may not be COVID-19 specific). The data used in the study^2^ came from an observational study^1^ in which all subjects from both treatment group and control group were hospitalized patients, including non-ICU patients and ICU patients. As a result, the genes identified can be classified as COVID-19 specific. These five critical genes are the smallest subset of genes reported ever in the literature. They describe nonlinear and competing interaction gene-gene relationships, different from pathways that have been widely studied in the literature. These five genes also form signature patterns as new biomarkers and classify COVID-19 patients into seven subgroups and heterogeneous populations.

The 100% accuracy proves the biological equivalences between COVID-19 status and the signature patterns and mathematically establishes the correspondences. Such a phenomenon is fundamentally meaningful to conduct further focused research on these five genes and those highly correlated genes to these five, leading to the second (third) generation of vaccines, antiviral drugs, and treatment methods.

The analytical method used in the study^2^ has been proven a powerful approach in the studies^13;14;15^ where breast cancer patients, colorectal cancer patients, and lung cancer patients are again classified into their respective groups with the highest accuracy (100%) among eleven different study cohorts with thousands of patients. These studies prove that the study method is a powerful and unique method that can be particularly useful for COVID-19 and cancer studies.

This paper is going to advance further the signature patterns found in the study^2^ using a different dataset generated by the study^1^ and compare the results from the study^2^ and this new work with a different dataset collected using different measurements. This paper is completely new in its conceptual framework in biological and mathematical equivalence compared with an earlier pure data analysis. This new study conducted a competing risk analysis using the max-linear logistic regression model to analyze 126 blood samples from COVID-19-positive and COVID-19-negative patients. The two groups are: the lab-confirmed COVID-19 hospitalized patients, the control is other disease types of hospitalized patients, including ICU cases. There are two types of datasets available. One type is TPM (Transcripts Per Million)^2^, while another type is EC (expected counts), which are analyzed in this paper. Both datasets led to competing COVID-19 risk classifiers derived from 19,472 genes and their differential expression values. The final classifier model only involves five critical genes, ABCB6, KIAA1614, MND1, SMG1, RIPK3, which led to 100% sensitivity and 100% specificity of the 126 samples. Given their 100% accuracy in predicting COVID-19 positive or negative status, these five genes can be critical in developing proper, focused, and accurate COVID-19 testing procedures, guiding the second-generation vaccine development, studying antiviral drugs and treatments. Furthermore, it is expected that these five genes can motivate numerous new COVID-19 researches.

Simultaneously observing the same set of five genes for two different datasets hasn’t been reported in published literature papers. In our opinion, those published genes by many other studies are more like at the surface level based on the analysis methods used, and the new set of five genes in this work is at the deep level or the root level. Many reported key genes are based on their individualized expression value changes and significance, i.e., not based on gene-gene interactions. As a result, treatments are palliative, and the disease can hardly be cured. The findings in our new research are based on nonlinear and competing risk factors interactions, which is an advanced gene-gene interaction mechanism. Our proposition is that the biomedical discovery of new variants of COVID-19 is only the surface-level of the virus (diseases). There are more profound, underlying “competing factors” of the virus that need to be studied. Metaphorically, an expert in hydraulic engineering finds a dam with cracks and treats them on the surface. However, the reservoir has an interconnected water dynamic below the surface that will further impact other points of the dam. As a result, it will cause new cracks unless there is a fundamental treatment solution with the entire structure in mind. Similarly, scientists may observe the variants (rather passive) and develop vaccines in response to the variants. However, they don’t understand the virus’s deeper advanced genomic architecture that will systematically cause other mutations. Traditional methods in statistics, machine learning, and AI are limited to understanding COVID-19 from surface-level observations. However, our innovative method has achieved significant results (100% accuracy) to identify and understand COVID-19 genomic signatures.

Among these five genes, KIAA1614 is an uncharacterized protein gene, according to genecards.org. This gene can be fundamental. SMG1 has the description: Nonsense Mediated mRNA Decay Associated PI3K Related Kinase. It can be essential for developing second-generation mRNA vaccines. According to the literature, Nonsense Mediated mRNA Decay plays a decisive role in monitoring and controlling protein changes. The mRNA gene SMG1 showed two different signs (+,-) in two formulas and didn’t appear in the third formula presented in Section 3, which suggests that we need three different types (both mRNA and non mRNA) of vaccines to cover the entire possible spectrum of COVID-19 variants. Our proven classifiers show that one particular vaccine may only be effective to one type of virus caused by one signature pattern and one subtype among three signature patterns and seven subtypes, which can explain the high percentages of breakthrough infections; see Sections 3.3 and 3.4 for detailed explanations.

Our findings can be used to develop precision test kits for testing COVID-19 and to evaluate the function and performance of already implemented vaccines, i.e., used as new antibody indexes. Interpretable and implementable formulas are given in the paper. After a COVID-19 case is confirmed, personalized treatments can be implemented. For example, increasing or decreasing levels of critical genes based on the identified COVID-19 subtype can be crucial to the patients’ recovery. Using the relationship determined in the findings, antiviral drugs can be developed. Mathematically, the new objective function of equation (4) is a mixture of combinatorial optimization and continuous optimization. It is a new type of classification benchmark which contains logistic regression, probit link, and Gumbel link as its special cases. It is expected that this new classification formula will motivate many research studies in statistics, computational mathematics, computer science, and many applied sciences. The findings can motivate many new types of research in COVID-19 studies and other studies, e.g., cancer studies. Many finished studies can re-start new looks using the new methodology.

The newly identified genes and their combinations may be used as new biomarkers. In our opinion, traditional methods (e.g., PCR, serology) are directly associated with the disease, i.e., they do not provide pathological characteristics; they are onefold indicators. On the other hand, the new classifier is a multifold indicator that can further divide the disease into subgroups (variants may be another word). In addition, the new classifier shows gene-gene interactions and advanced (or root) structures.

This work has verified that when all component classifiers work, the patients are ICU patients, which definitely points out the advanced genomic structure of COVID-19 disease.

In the literature and the current practice, tremendous efforts have been made to study COVID-19 genomic sequences, variants and their impacts, and vaccine effectiveness. However, the progress on the pathological causes of COVID-19 and the functional effects of genes is still limited. In terms of computational medicine, our new work is the first to accurately define the functional effects of five critical genes and lead to the mathematical and biological equivalence between five genes and COVID-19 status. Furthermore, this paper introduces an advanced machine learning algorithm that identifies five essential genes, which further determine three genomic signature patterns and seven subtypes of COVID-19 with 100% accuracy. The final classifiers are expressed by explicit mathematical equations which are interpretable and can guide medical practice. In addition, new graphical diagnostic tools are introduced. Besides the strike advance of studying genomic signature patterns of COVID-19, our work also sheds new light on computational medicine, genetic studies, informatics, algorithm and machine learning, and statistics. The rest of the paper is organized as follows. In Section 2, the methodology is briefly summarized. Section 3 presents the data, the derived competing classifiers, and the interpretations. In Section 4, discussions on the findings, future directions and limitations are discussed.

## 2 Methodology

This methodological section briefly introduces max-linear competing factor classifiers for self-contained due to different data structures used in this work compared with other researches whose details for data structures can be found in the papers^2;13;14;15^. For continuous responses, we refer to max-linear computing factor models and max-linear regressions with penalization in the literature^16;17^.

Suppose *Y*_*i*_ is the *i*th individual patient’s COVID-19 status (*Y*_*i*_ = 0 as not infected, *Y*_*i*_ = 1 for infected) and 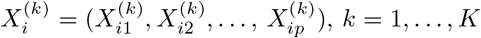, being the gene expression values with *p* = 19472 genes in this study. Here *k* stands for the *k*th type of gene expression levels drawn based on *K* different biological sampling methodologies. Note that most published work set *K* = 1, and hence the supercript (*k*) can be dropped from the predictors. In this research paper, *K* = 2. Using a logit link (or probit link, Gumbel link), we can model the risk probability 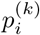 of the *i*th person’s infection status as:

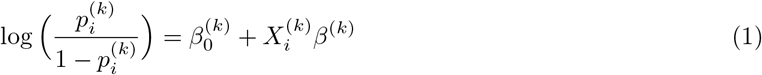

or alternatively, we write

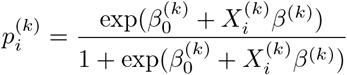

where 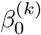 is an intercept, 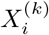 is a 1 *× p* observed vector, and *β*^(*k*)^ is a *p ×* 1 coefficient vector which characterizes the contribution of each predictor (gene in this study) to the risk.

Considering there have been several variants of SARS-COV-2 and multiple symptoms (subtypes) of COVID-19 diseases, it is natural to assume that the genomic structures of all subtypes can be different. Suppose that all subtypes of COVID-19 diseases may be related to *G* groups of genes

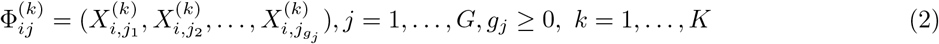

where *i* is the *i*th individual in the sample, *g*_*j*_ is the number of genes in *j*th group. Note here, we do not use the widely used gene pathways in our newly developed machine learning algorithm. It is possible that blind pursuit of gene pathways may lead to wrong directions and lose chances of finding the scientific truth. Instead, our methodological approach will automatically find the smallest numbers of *G* and *g*_*j*_ and to reach a 100% accuracy, and as a result, better or best interpretations can be achieved.

The competing (risk) factor classifier is defined as

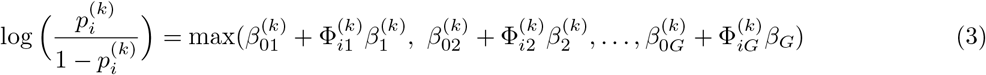

where 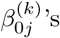s are intercep ts, 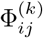 is a 1*×g*_*j*_ observed vector, 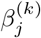 is a *g*_*j*_ *×*1 coefficient vector which characterizes the contribution of each predictor in the *j*th group to the risk.

### Remark 1.

*In (3)*, 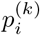 *is mainly related to the largest component* 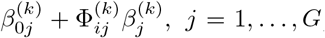, *i*.*e*., *all components compete to take the most significant effect*.

### Remark 2.

*Taking* 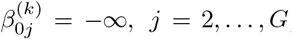, *(3) is reduced to the classical logistic regression, i*.*e*., *the classical logistic regression is a special case of the new classifier. Compared with blackbox machine learning methods (e*.*g*., *random forest, deep learning (convolution) neural network (DNN, CNN)) and regression tree methods, (3) shows clear patterns. Each competing risk factor forms a signature with the selected genes. The number of factors corresponds to the number of signatures, i*.*e*., *G. This model can be regarded as a bridge between linear models and more advanced (blackbox) machine learning methods. However, (3) remains the desired properties of interpretability, computability, predictability, and stability. Note that this remark is the same as Remark 1*^*15*^.

In practice, we have to choose a threshold probability value to decide a patient’s class label. Following the general trend in the literature, we set the threshold to be 0.5. As such, if 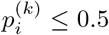, the *i*th individual is classified as disease free, otherwise the individual is classified to have the disease.

With the above established notations and the idea of quotient correlation coefficient^18^, Zhang (2021)^15^ introduces a new machine learning classifier, smallest subset and smallest number of signatures (S4), for *K* = 1. We extend the S4 classifier from *K* = 1 to *K* = 2 as follows.

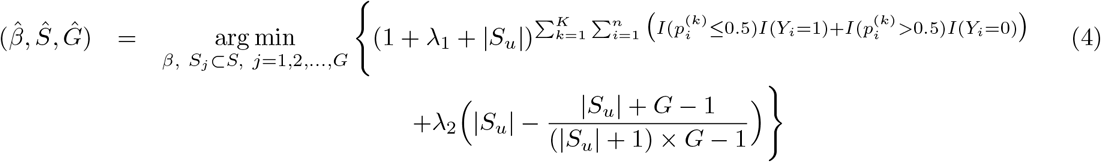

where *I*(*·*) is an indicative function, 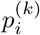 is defined in Equation (3), *S* = *{*1, 2, …, 19472*}* is the index set of all genes, *S*_*j*_ = *{j*_*j*1_, …, *j*_*j,gj*_ *}, j* = 1, …, *G* are index sets corresponding to (2), *S*_*u*_ is the union of *{S*_*j*_, *j* = 1, …, *G}*, |*S*_*u*_| is the number of elements in *S*_*u*_, *λ*_1_ *≥* 0 and *λ*_2_ *≥* 0 are penalty parameters, and *Ŝ* = *{j*_*j*1_, …, *j*_*j,gj*_, *j* = 1, …, *Ĝ}* and *Ĝ* are the final gene set selected in the final classifiers and the number of final signatures.

### Remark 3.

*The case of K* = 1 *corresponds to the classifier introduced in Zhang (2021)* ^*15*^. *The case of K* = 1 *and λ*_2_ = 0 *corresponds to the classifier introduced in Zhang (2021)* ^*2*^.

### Remark 4.

*A perfect classifier (100% sensitivity and 100% specificity) will have* 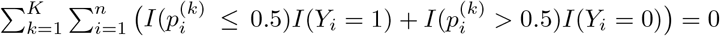 *in Equation (4), which is the case in our study*.

The following Proposition 2.1 mathematically guarantees the existence of desired solutions. The proof follows the lines in the work^15^ by extending *K* = 1 to *K* = 2.

### Proposition 2.1.

*Suppose the smallest number that* 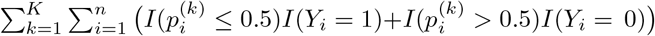 *can reach is m. Then for suitable choices of λ*_1_ *with λ*_1_ + |*S*_*u*_| *>* 0 *and λ*_2_ *≥* 0, *the new classifier S4 will lead to the smallest* |*S*_*u*_| *and the smallest number of G such that* 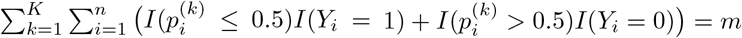.

## 3 Data Descriptions, Results and Interpretations

### 3.1 The data

The COVID-19 data to be analyzed is publicly available as GSE157103: Large-scale Multi-omic Analysis of COVID-19 Severity^1^, Public on August 29, 2020. The experiment type is “Expression profiling by high throughput sequencing.” One hundred twenty-six samples were analyzed in total, with 100 COVID-19 patients and 26 non-COVID-19. There are two types of datasets available. One type is TPM (Transcripts Per Million), while another type is EC (expected counts). The prior analysis outcome of TPM data was reported in Zhang (2021)^2^. This new study targets EC data and makes comparisons to TPM data. We note that among 100 COVID-19 patients, 50 are ICU patients, and 50 are non-ICU hospitalized patients. Among 26 COVID-19 free patients, 16 of them are ICU patients with other types (non-COVID-19) of disease, and 10 of them are non-ICU patients with other types of disease.

### 3.2 The competing factor classifiers and their resulting risk probabilities

Solving the optimization problem (4) among 19472 genes with *K* = 2 (*k* = 1 for TPM data, and *k* = 0 for EC data), we obtain the following classifiers with five critical genes (ABCB6 (ATP Binding Cassette Subfamily B Member 6 - Langereis Blood Group), KIAA1614 (Uncharacterized Protein), MND1 (Meiotic Nuclear Divisions 1), SMG1 (Nonsense Mediated mRNA Decay Associated PI3K Related Kinase), RIPK3 (Receptor Interacting Serine/Threonine Kinase 3)):

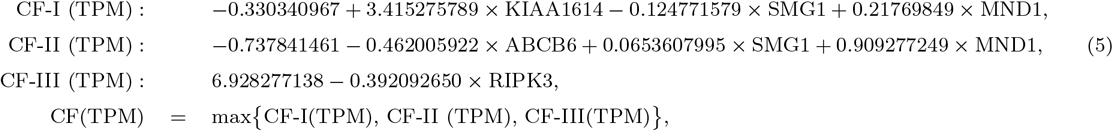

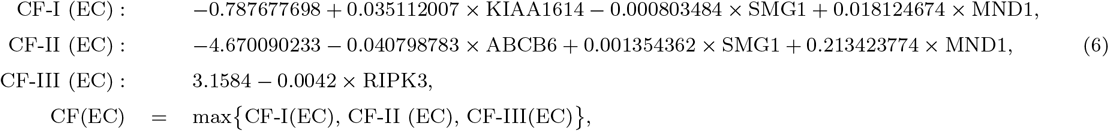

where Equations (5) are for TPM data which were first reported in Zhang (2021)^2^, while Equations (6) are for EC data. In all equations, (TPM) stands for data being TPM, and (EC) means data are expected counts. In Equations (5), CF-I (TPM) is the first component classifier derived from TPM data and three critical genes KIAA1614, SMG1, and MND1. CF-II (TPM) is the second component classifier derived from TPM data and three critical genes ABCB6, SMG1, and MND1. CF-III (TPM) is the third component classifier derived from TPM data using one gene RIPK3 alone. CF (TPM) is the final combined classifier with competing component classifiers (signatures) of CF-I (TPM), CF-II (TPM), and CF-III (TPM). Other competing classifiers for EC are similarly defined.

The risk probabilities of each of three component classifiers based on TPM are

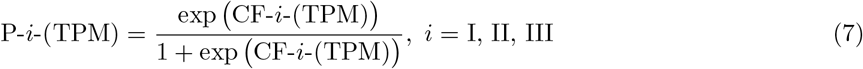

and the risk probabilities based on all three component classifiers together are

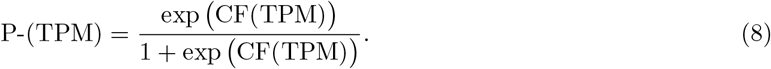

Similarly, the risk probabilities calculated from EC are

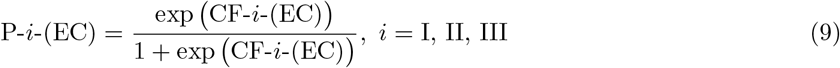

and the risk probabilities based on all three component classifiers together are

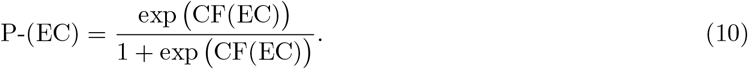

Tables 1 and 2 present some patients’ gene expression values, competing classifier factors, predicted probabilities defined in equations (5-10). For full tables, we refer to tables in a supplementary Excel file.

**Table 1:**
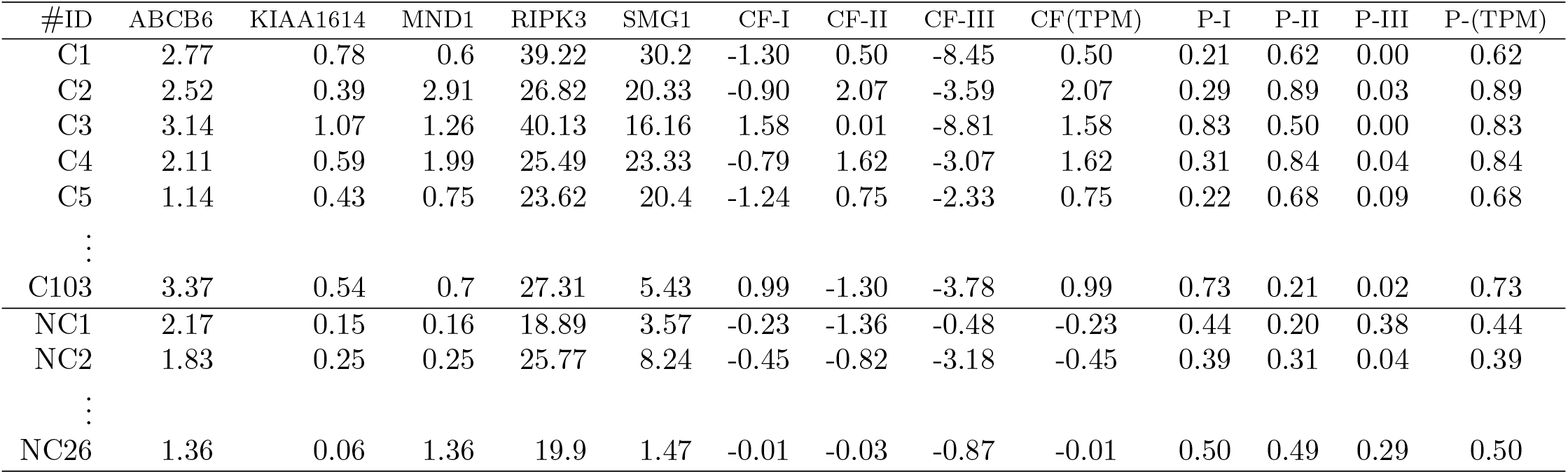
TPM data: Expression values of the five critical genes, competing classifier factors, predicted probabilities. C1,…, C103 are hospitalized COVID-19 patient IDs, and NC1,…, NC26 are COVID-19 free patient (also hospitalized) IDs.

**Table 2:** EC data: Expression values of the five critical genes, competing classifier factors, predicted probabilities. C1,…, C103 are hospitalized COVID-19 patient IDs, and NC1,…, NC26 are COVID-19 free patient (also hospitalized) IDs.

| #ID | ABCB6 | KIAA1614 | MND1 | RIPK3 | SMG1 | CF-I | CF-II | CF-III | CF(EC) | P-I | P-II | P-III | P-(EC) |
| --- | --- | --- | --- | --- | --- | --- | --- | --- | --- | --- | --- | --- | --- |
| C1 | 143 | 141.2 | 8 | 1217 | 8951.6 | -2.88 | 3.33 | -0.47 | 3.33 | 0.05 | 0.97 | 0.39 | 0.97 |
| C2 | 82 | 46.13 | 25 | 539 | 3913.32 | -1.86 | 2.62 | 0.21 | 2.62 | 0.13 | 0.93 | 0.55 | 0.93 |
| C3 | 159 | 195.76 | 17 | 1250 | 4803.49 | 2.53 | -1.02 | -0.50 | 2.53 | 0.93 | 0.26 | 0.38 | 0.93 |
| C4 | 92 | 92.4 | 23 | 685 | 6001.07 | -1.95 | 4.61 | 0.07 | 4.61 | 0.12 | 0.99 | 0.52 | 0.99 |
| C5 | 58 | 78.36 | 10 | 734 | 6085.32 | -2.74 | 3.34 | 0.02 | 3.34 | 0.06 | 0.97 | 0.50 | 0.97 |
| ⋮ |  |  |  |  |  |  |  |  |  |  |  |  |  |
| C103 | 290 | 169.66 | 16 | 1451 | 2772.87 | 3.23 | -9.33 | -0.70 | 3.23 | 0.96 | 0.00 | 0.33 | 0.96 |
| NC01 | 151 | 37.54 | 3 | 826 | 1479.41 | -0.60 | -8.19 | -0.07 | -0.07 | 0.35 | 0.00 | 0.48 | 0.48 |
| NC02 | 140 | 67.34 | 5 | 1199 | 3688.33 | -1.30 | -4.32 | -0.45 | -0.45 | 0.21 | 0.01 | 0.39 | 0.39 |
| ⋮ |  |  |  |  |  |  |  |  |  |  |  |  |  |
| NC26 | 126 | 22 | 35 | 1168 | 814.16 | -0.04 | -1.24 | -0.42 | -0.04 | 0.49 | 0.22 | 0.40 | 0.49 |

Figures 1 and 2 present critical gene expression levels and risk probabilities corresponding to different measurement scales and different component competing factor classifiers. From Figures 1 and 2, we can see clear patterns of how the patients are classified and how they are correlated with individual genes. For example, some patients can be classified by one gene RIPK3 (the right panels in the figures), while some patients are classified by the combined effects of linear combinations of three genes (the left and middle panels). As a result, these observations justify the existence of three genomic signature patterns, i.e., the three competing classifiers, of COVID-19.

**Figure 1:**
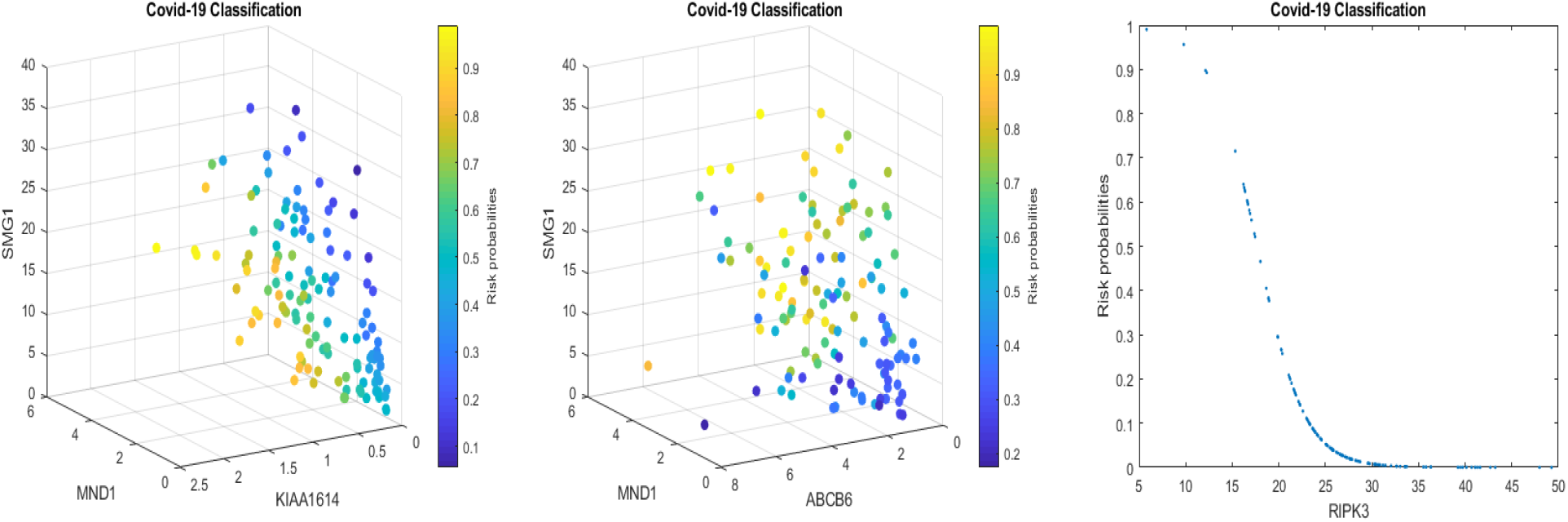
COVID-19 classifiers using TPM data: Visualization of gene-gene relationship and gene-risk probabilities. Note that 0.5 is the probability threshold.

**Figure 2:**
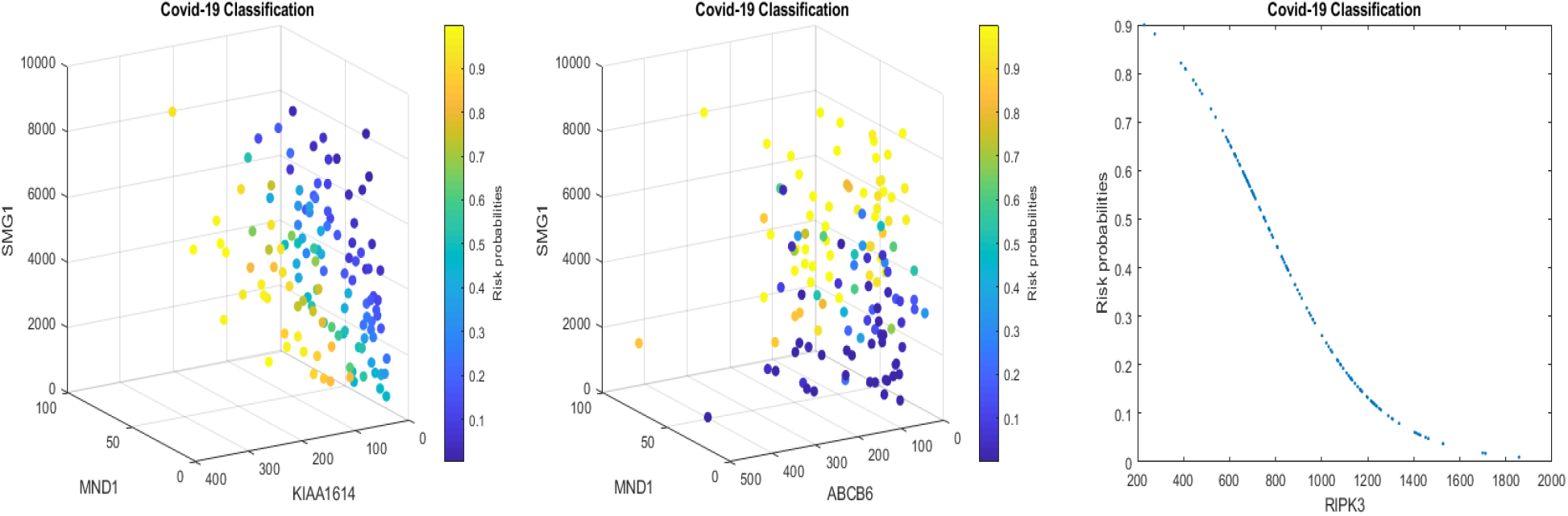
COVID-19 classifiers using EC data: Visualization of gene-gene relationship and gene-risk probabilities. Note that 0.5 is the probability threshold.

We can also see similar patterns between Figures 1 and 2. This phenomenon is mainly due to that the component genes and signs of coefficients in the classifiers CF-*i*-(TPM) and CF-*i*-(EC) are the same. In addition, the linear correlation coefficients between TMP and EC data for genes ABCB6, KIAA1614, MND1, RIPK3, SMG1, are 0.87, 0.94, 0.95, 0.68, 0.93, respectively, which supports the pattern similarity.

### 3.3 The combination effects and the competing factor effects

The same signs of coefficients in the classifiers CF-*i*-(TPM) and CF-*i*-(EC) reveal that these classifiers are robust to nonlinear transformations and the five genes are critical and COVID-19 specific (recall that the patients in the control group are also hospitalized patients with 16 of them being ICU patients.)

The pairwise correlation coefficients among these five genes are presented in Table 3. The table tells that TPM data and EC data show different gene-gene correlations. Even so, they still lead to the same accuracy (100%). As a result, they can be used as cross-validation of the proposed S4 model (4).

**Table 3:** Pairwise correlation coefficients: The upper triangle is for TPM data, and the lower triangle is for EC data.

|  | ABCB6 | KIAA1614 | MND1 | RIPK3 | SMG1 |
| --- | --- | --- | --- | --- | --- |
| ABCB6 | – | 0.6931 | 0.3448 | -0.1204 | 0.2138 |
| KIAA1614 | 0.5209 | – | 0.1609 | 0.1688 | 0.5948 |
| MND1 | 0.4009 | 0.1163 | – | -0.1328 | 0.1276 |
| RIPK3 | -0.0727 | -0.1608 | -0.0769 | – | 0.2293 |
| SMG1 | -0.0955 | 0.4681 | -0.0137 | -0.1762 | – |

From Equations (5) and (6), we can see that increasing ABCB6 and RIPK3 expression levels (TPM) will benefit the patients while decreasing KIAA1614 and MND1 expression levels will help the patients. However, from Table 3, we see that ABCB6 is positively correlated with KIAA1614 and MND1, and then an increase of ABCB6 expression level may result in an increase of MND1 expression level and KIAA1614 expression level which increases the COVID-19 competing risk CF-I. As a result, any efficient treatments of COVID-19 must consider all the five genes together.

Note that RIPK3 does not appear in the classifiers CF-I and CF-II, and the signs of SMG1 in CF-I and CF-II are different. As a result, a vaccine/antiviral drug/therapy which is based on the function of SMG1 (an mRNA gene) in the CF-I (CF-II) may benefit the patients classified in the CF-I (CF-II). However, it may not have any effects on the patients in the CF-III. Furthermore, it may make the status of patients in the CF-II (CF-I) worse. Analogously, a vaccine/antiviral drug/therapy which is based on the function of RIPK3 (CF-III) may not be effective to the patients caused by the effects of CF-I and CF-II. These observations reveal that we may need at least three different types of vaccines against COVID-19 subtypes (variants).

Note that these five critical genes have not been reported in any literature papers except Zhang (2021)^2^. They are not from any single gene pathway. Analogously, their functions may be described as a basketball team’s player combination effects. First, in a basketball team, there are five positions: center (C), power forward (PF), small forward (SF), point guard (PG), shooting guard (SG). A combination of ABCB6-MND1-SMG1 (KIAA1614-MND1-SMG1) may be described as a driving force of a powerful PF-C-PG (SF-C-PG) combination in scoring, and RIPK3 is like a powerful SG. Second, the expression levels are comparable to the playing time of the players and their roles in the rotations competing against different opponents and their playing combinations. Third, the driving forces of winning games can be either one or two or all of the three combinations.

Note that the correlation coefficients among the five genes calculated from TPM (upper triangle) and EC (lower triangle) in Table 3 are different. This phenomenon can be explained by the nonlinear relationship between TPM and EC, within TPM and EC, and heterogeneous populations among patients, which is a perfect scenario for the proposed model in Section 2. Also, note that due to 100% accuracy, metrics such as ROC, Recall, and Precision are also with 100% accuracy.

### 3.4 The existence of subtypes

In Section 3.2, we saw that each signature could be used as a classifier given its 100% specificity. However, from Figures 1 and 2, we see that some patients can only be classified by one particular signature classifier; some patients can only be classified by two competing classifiers, and some patients can be classified by either of the three competing classifiers. This observation shows that COVID-19 patients are heterogeneous, and their COVID-19 status can be further classified into subtypes using the classifier combinations.

Figure 3 uses Venn diagrams to plot classified seven subtypes of 100 COVID-19 patients. In the figure, I-II means both CF-I and CF-II lead to the correct classification. All other combinations are interpreted similarly.

**Figure 3:**
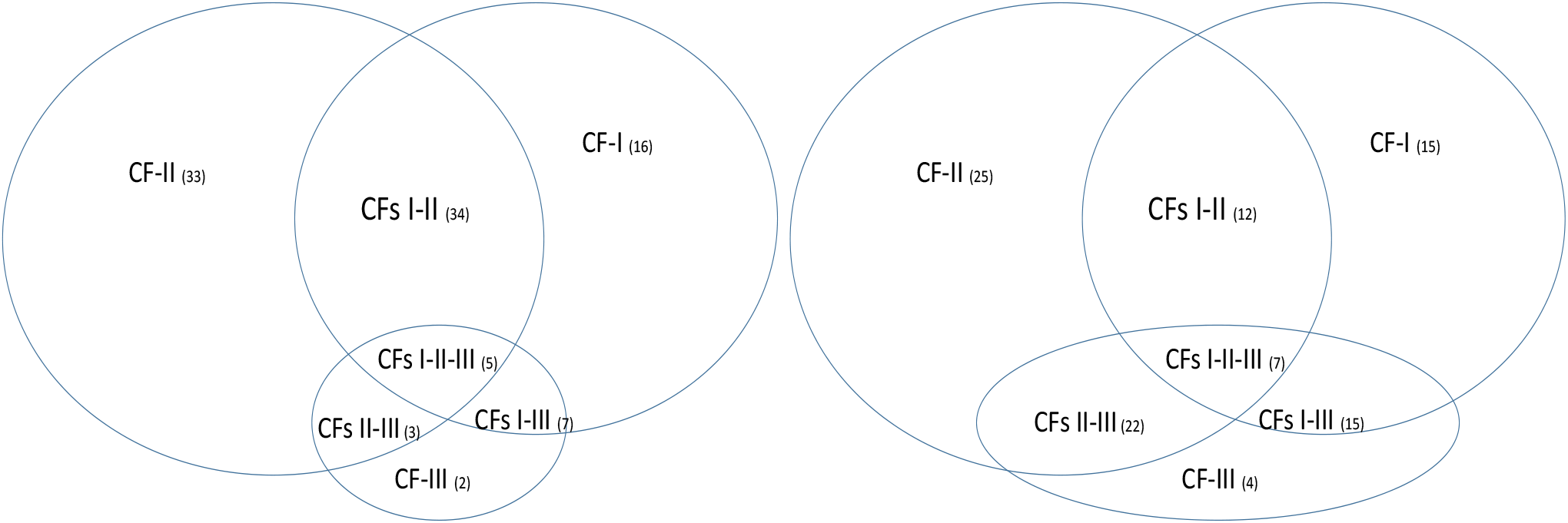
Venn diagrams of COVID-19 subtypes. The left panel is based on TPM data, and the right panel is based on expected counts data.

We first state that the more intersections of subareas, the more complicated the disease in a Venn diagram. For example, in the right panel of the figure, we see that seven patients satisfy the classification conditions of all three competing classifiers. It turns out all of these seven patients are ICU COVID-19 patients. Using the right panel as an example, we identify the ICU patients have the following distribution: CF-I (6), CF-II (8), CF-III (2), CFs-I-II (6), CFs-I-III (12), CFs-II-III (9), and CFs-I-II-III (7), and find that the disease severity (ICU) is positively correlated with the number of classifiers used. Based on this observation, we can conclude vaccines can benefit patients even if a particular type of vaccine only works for one signature pattern related to COVID-19 subtypes. On the other hand, if one particular type of vaccine can protect a particular COVID-19 subtype (or SARS-COV-2 virus), this vaccine may not be effective for other COVID-19 subtypes (or SARS-COV-2 viruses.) As a result, a fully vaccinated individual still has the risk of being infected. Furthermore, a recovered individual from an infected COVID-19 illness can get breakthrough infections again with other COVID-19 subtypes. Two recent papers^19;20^ report concerns of infection-enhancing anti-SARS-CoV-2 antibodies based on lab experiments. This phenomenon may be explained by our new findings due to the existence of three genomic signature patterns and seven subtypes of COVID-19. Taking the SMG1 gene as an example, an increase (or a decrease) of SMG1 expression levels is good for one signature pattern of COVID-19 but will be bad for another signature pattern.

Note that the left panel and the right panel classify some patients into different subtypes. This phenomenon can be explained. In Section 3.2, we calculated the linear correlation coefficients between TPM and EC for each gene and saw there are nonlinear relationships, which lead to different classifications, but still 100% accurate. Given that both TPM and EC lead to 100% accuracy, it is safe to say that the identified five genes can be truly critical in studying COVID-19.

The identified subtype information opens a new research direction, new drug developments, and new refined personalized therapies. For example, in the diagnosis stage, medical doctors can use the final model to predict their patient’s COVID-19 status by calculating the risk and determine which of those seven groups the patient belongs to. In the treatment stage, those signature patterns can be used to study the effectiveness of drugs and treatments, i.e., conduct clinical trials based on classified groups. In the drug development stage, pharmaceutical companies can use the findings of critical genes to study new drugs. Finally, it can be hoped that mRNA-based therapies can be introduced using the critical genes’ information in the therapy stage.

### 3.5 A conceptual framework

From Section 3.2, we see that COVID-19 patients have higher combination expression values, COVID-19 free patients have lower combination expression values. With 100% sensitivity and 100% specificity, the competing classifiers derived in Section 3.2 have built a biological equivalence to COVID-19 status. Equations (5) and (6) together with 100% sensitivity and 100% specificity reveal the hyperplanes formed by five genes and their derived classifiers partition a five-dimension space into two subspaces (COVID-19 and COVID-19 free) in which there is a mathematical equivalence between COVID-19 and COVID-19 free. Here the logic is from the fact of 100% sensitivity and 100% specificity, i.e., if A implies B, then not B implies not A; and if not A implies not B, then B implies A.

In the Introduction, we used hydraulic engineering of a reservoir dam with cracks to describe COVID-19 variants metaphorically. In Figure 4, we use the five genes identified in Section 3.2 and one additional gene CDC6 (to be discussed in Section 4) to describe a conceptual framework for COVID-19 disease and variants formation dynamics.

**Figure 4:**
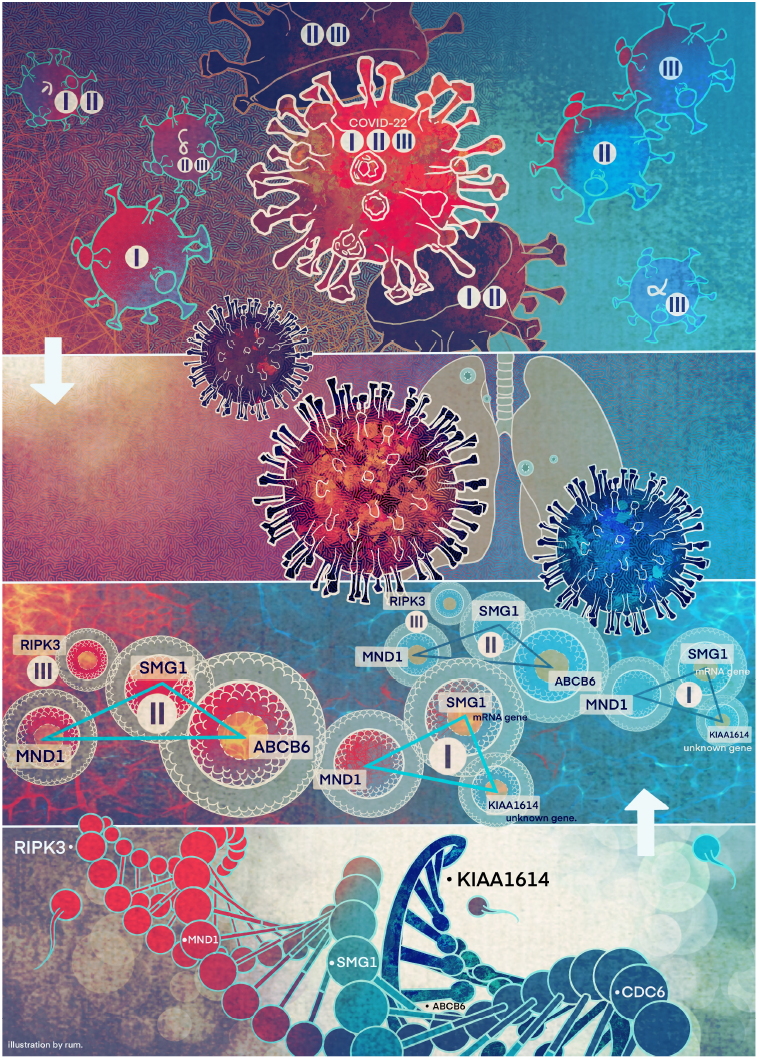
COVID-19 formation: A conceptual visualization of gene-gene relationship. The author designed the concept and flowchart. Jing Zhang completed drawing the figure.

In Figure 4, there are four layers. The first (top) layer stands for SARS-COV-2 viruses enter a human’s interior body. The second layer shows the lung will be affected. The third layer describes gene-gene interaction signature patterns of COVID-19. The bottom layer is a conceptual illustration of a human genome sequence with the five critical genes placed on the sequence. In the figure, we use triangles to represent signature patterns (competing factor classifiers) with the genes on the nodes and the classifier number inside the center. RIPK3 is on its own as an absorbed triangle. There are two arrows to indicate the cause dynamics. With the triangles or RIPK3, the larger the triangle or the shape of RIPK3, the higher the severity of the COVID-19. Our conceptual idea is that after being infected with COVID-19 (top-down direction), the virus triggers the signature patterns to function, i.e., making the triangles (the shape) larger. However, simultaneously, the human’s immune system starts to function (bottom-up direction), and the vaccine also starts to function; therefore, the areas of triangles (shape) can be reduced, or the edges of the triangles can be broken, i.e., two ways of fighting against the virus. Depending on which direction (infection or killing the virus) is more effective, an infected individual may be fully recovered from COVID-19 disease or going to be much severer COVID-19 symptoms.

On the other hand, the genomic signature patterns of a COVID-19 patient represent the advanced (deep level) gene-gene interactions. A change of the size of the triangle may trigger new variants to form and transmit to other individuals, i.e., an analog to the hydraulic engineering example in Introduction.

## 4 Discussions

As discussed in Sections 1 and 3, the three signature patterns and seven subtypes maintain the most important biological informatics of COVID-19. This set of genes is the only set that leads to 100% accuracy for hospitalized COVID-19 patients, including ICU patients. Unless a different new discovery of other advanced structures of genes other than these five genes will be obtained, and such a new discovery (if exists) can fully explain the three signature patterns and seven subtypes discussed in this paper, these five critical genes and their derived three signature patterns and seven subtypes remain the most informative findings.

When a model is fitted to the whole dataset and leads to 100% accuracy, it will uniformly work for partitioned data as long as the partition is balanced to all heterogeneous subgroups. This is the case in our analysis. On the other hand, we haven’t seen any published papers that used the “standard” procedure to lead to accurate prediction.

We realized readers would ask about the model overfitting and robustness. Please note that our model is fitted to two different datasets and reached the highest accuracy of 100%. Each dataset has its heterogeneous patterns (subgroups). Datasets are measured at different scales. It’s hard for the existing models to simultaneously fit such datasets and get satisfactory accuracy, not even to mention the interpretability of the fitted models. Using two such datasets naturally serves as cross-validation and robust checking. It turns out our new approach is robust.

A 100% accuracy may be thought of as “too good to be true” in many scenarios. However, “too good to be true” may also be dangerous to use to guide science discovery and innovation. In many applied sciences, the truths can be simple but not straightforward. Complicating or aggravating the problem can mask the nature of the problem. Blindly insisting on “too good to be true” may miss ample opportunities of finding the truth. In contrast, we know it is hard to see the forest through the trees.

One may argue that the dataset we used in this analysis is not large enough as it has only 126 samples with 19472 predictors. It is, of course, preferred to have a large dataset. Nevertheless, we argue that the conclusions and inferences are trustful with 100% accuracy on two different datasets that show nonlinear and heterogeneous relationships.

On the other hand, if an approach can not gain a satisfactory performance with a small dataset, applying it to a large dataset can be a wrong strategy as it may lead to wrong or suboptimal conclusions.

A natural question is whether or not the 100% accuracy is by chance. Note that each of our component competing classifiers has reached 100% specificity, which may be true with a probability smaller than 1*/*2^26^ = 1.0 *×* 10^*-*8^ by chance. In addition, when all three signature patterns are satisfied, all classified patients are lab-confirmed ICU patients, which indicates it can not be by chance.

We note that multiplying a constant to Equations (5-6) will not change the classification results and the shapes in Figures 1-2 with a 100% accuracy being achieved. However, the color strengths will be changed. Such a phenomenon justifies the use of signatures to describe the advanced gene-gene interaction structures. Using this idea, we can express (6) into the following equivalent forms:

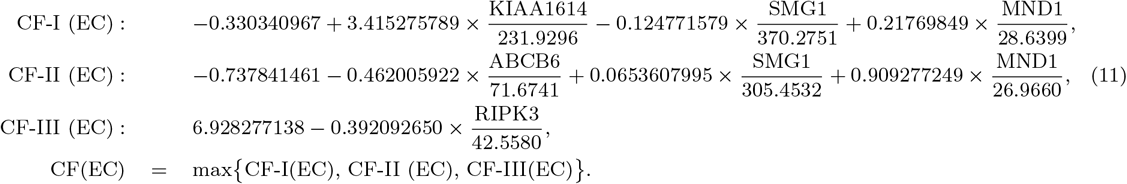

Equations (11), after applying scale changes, share similar signatures as in Equations (5). Considering the nonlinear relationships between TPM data and EC data, this observation again proves our proposed competing factor classifiers are robust.

We want to suggest that the discovered COVID-19 variants (alpha, beta, delta, lambda, mu, omicron etc.) may be connected to different signature strengths in our discovered signature patterns. Mathematically, hyperplanes in geometry formed by Equations (5) and (11) contain a subspace which can be further partitioned into subspaces. Our hypothesis is that these variants may fall into separable subspaces. After obtaining new data with variants information, this hypothesis can be tested, or additional genes may be involved. For example, in a breast cancer study^13^, triple-negative breast cancer was 100% accurately separated from other types of breast cancer using three genes identified by the S4 classifier. Furthermore, the discovered genomic signature patterns and COVID-19 subtypes are intrinsic no matter what variants have been identified or will be identified. Given our proven mathematical and biological equivalences, if these innate signature patterns and subtypes cannot be treated and fully studied now, they will cause trouble in the future again. In addition, with available data related to various variants, our study approach may be able to reveal the causes of higher transmission or mortality of specific variants.

Finally, in our analyses, we also found the gene CDC6 (cell division cycle 6) can be informative. Its combination to ZNF282 (Zinc Finger Protein 282) can lead to 97.62% accuracy (98% sensitivity, 96.15% specificity), and its combination to both ZNF282 and CEP72 (Centrosomal Protein 72) can lead to 98.41% accuracy (99% sensitivity, 96.15% specificity). We found the high expression level of CDC6 increases risk while the high expression levels of ZNF282 and CEP72 decrease risk. Although they didn’t lead to 100% accuracy as those five identified in Section 3 did, such a performance is already better than other published gene combinations. From the literature, the gene CDC6 is a protein essential for the initiation of DNA replication, while ZNF282 is known to bind U5RE (U5 repressive element) of HLTV-1 (human T cell leukemia virus type 1) with a repressive effect but little is known of its expression and function otherwise. The gene CEP72 coded protein is localized to the centrosome, a non-membraneous organelle that functions as the major microtubule-organizing center in animal cells. Zhang^21^ hypothesizes that CDC6 is a protein essential for the initiation of RNA replication of COVID-19.

## Acknowledgments

The author thanks anonymous editors and anonymous reviewers for their valuable comments and suggestions for the first two versions of the paper. The author also thanks the helpful information, discussions and assistance from Cynthia Czajkowski, Thomas Friedrich, Richard Smith, Jun Yan, Lei Qin, Hanwen Su, Xuming He, Ariel Jaitovich, Joseph Balnis, Joshua Coon, Xihong Lin, Jianqing Fan, Zhiliang Ying, Jing Qin, Dean Follmann, Yuqing Xu, Qiurong Cui, Hao Yang Teng, Xiahai Zhuang, Jerry Yin, Wei Xu, Ariel Qihui Tao, Jing Zhang and many other people who discussed the project with the author and encouraged author to put effort in this urgently needed study. The partial support from NSF-DMS2012298 is acknowledged.

## Supplementary materials

Real data and computer outputs are in a supplementary file available online and submitted together with this paper. A Matlab ^®^demo code for solving *A* in Equation (4) (*λ*_2_ = 0) is also available.

## Data availability

The datasets are publicly available. The data links are stated in Section Data Description.

## Limitation statements

Data used in this study are from hospitalized patients’ blood samples. We are not sure whether or not the identified genes work for data sampled from those non-hospitalized COVID-19 patients or asymptomatic patients. Solving optimization problems (4) involves combinatorial optimization, integer programming, and continuous programming. The computational complexity is extremely high, and we haven’t figured out how to define the complexity. We used an extensive Monte Carlo search method to find the best solution. However, we cannot guarantee whether additional sets of genes can also be the optimal solutions even if our finding of five genes is already the best (smallest) subset genes with 100% accuracy in the study of COVID-19. Although we have established the mathematical and biological equivalences, we cannot tell our findings are the causes or the results. Although we have identified functional effects by gene-gene interactions and gene-subtype interactions of the five genes, we haven’t identified how gene-gene interacts each other and their causal directions. We are working in this direction. Finally, due to the lack of available new blood sampled data for new variants, it’s difficult to infer the risks of variants.

